# Mapping plant-scale variation in crop physiological traits and water fluxes

**DOI:** 10.1101/2025.02.21.639464

**Authors:** Robert S. Caine, Peter M. Berry, Kate E. Storer, Holly L. Croft

## Abstract

Nitrogen (N) is a vital plant element, affecting plant physiological processes, carbon and water fluxes and ultimately crop yields. However, N uptake by crops can vary over fine spatiotemporal scales, and optimising the application of N-fertiliser to maximise crop performance is challenging. To investigate the potential of spatially mapping the impact of N fertiliser application on crop physiological performance and yield, we leverage both optical and thermal data sampled from drone platforms and ground-level leaf measurements, across a range of different N, Sulphur (S) and sucrose treatments in winter wheat. Using leaf level hyperspectral reflectance data, leaf chlorophyll content was accurately modelled across fertiliser treatments via partial least squares regression (PLSR; *R^2^*= 0.93, *P* < 0.001). Leaf photosynthetic capacity (*V*_cmax_) exhibited a strong linear relationship with leaf chlorophyll (*R^2^* = 0.77; *P* < 0.001). Using drone-acquired MERIS terrestrial chlorophyll index (MTCI) values as a proxy for leaf chlorophyll (*R^2^* = 0.76; *P* < 0.001), *V*_cmax_ was spatially mapped at the centimetre-scale. Thermal drone and ground measurements demonstrated that N application leads to cooler leaf temperatures, which led to a strong relationship with ground-measured leaf stomatal conductance (*R^2^*= 0.6; *P* < 0.01). Final grain yield was most accurately predicted by optical reflectance (MTCI, *R^2^* = 0.94; *P* < 0.001). Precise retrieval of leaf-level crop performance indicators from drones establishes significant potential for optimising fertiliser application, to reduce environmental costs and improve yields.

## 1. Introduction

Nitrogen (N) is an essential plant macronutrient, affecting plant growth, productivity and a swathe of metabolic pathways (Caine et al. 2024; Guo et al. 2007). Since the middle of the 20^th^ century, the global use of synthetic nitrogen-based fertilisers to boost crop yields has increased 9-fold, from 11.6 Tg N yr^−1^ in 1961 to 109.1 Tg N yr^−1^ in 2017 (Ludemann et al. 2024; Yulong et al. 2021). This increased application of fertiliser has played a key role in the 250% growth in crop yields over the same timescale (Blomqvist et al. 2020). However, there are financial, societal and environmental costs associated with both the production and application of N-based fertiliser (Alexander et al. 2023; Wang et al. 2018). From an environmental position, fertiliser is often applied excessively; which can result in unwanted losses of N to the environment through emissions, volatilization and or leaching (Ladha et al. 2020; Plett et al. 2020). The rise in nitrogen fertiliser use has led to increased emissions of N_2_O, a gas nearly 300 times more potent than CO_2_ over a 100 year period, and a major contributor to climate change (Griffis et al. 2017). There is therefore a pressing need to optimise N-fertiliser application, whilst simultaneously maintaining or sustainably enhancing yields.

Early detection of deficiencies in crop N is critical for boosting N fertiliser-use efficiency and yields (Plett et al. 2020). Optimising the balance between meeting the physiological requirements of crops, whilst simultaneously minimising losses to the environment, involves the optimal rate, placement and timing of fertiliser application (Johnston and Bruulsema 2014), which requires targeted and dynamic monitoring of crop nutrient status (Berger et al. 2020). Different remote sensing techniques provide an opportunity to harness information content according to different mechanistic processes associated with crop stress to map within-field spatial variations in crop performance at user-defined temporal intervals. When first N becomes limiting, photosynthesis rates reduce and plants adjust their stomata to limit water loss, leading to decreases in latent-heat (LE) fluxes (McNellis et al. 2024; Plett et al. 2020). These N stress-induced changes in LE can be sampled using thermal remote sensing, as canopy temperature is inversely related to evaporative cooling as plants transpire (Berger et al. 2022). In addition to reduced LE, lower N also reduces photosynthetic capacity leading to reduced CO_2_ uptake and H_2_O release (Caine et al. 2024). Leaf functional traits such as the maximum rate of rubisco carboxylation (*V*_cmax_), leaf chlorophyll content and leaf N content are typically lower when N is insufficient (Croft et al. 2017). These changes in biochemical and physiological traits usually occur over timescales of days to weeks and can be detected using optical sensors based on the ground or via drone platforms which collect multispectral or hyperspectral data (Xie and Yang 2020). The accumulation of higher N input over time leads to increases in leaf area index (LAI) and larger biomass (Caine et al. 2024), which also affects crop energy balance as less soil understory is exposed, leading to lower sensible heat flux (H) (Yang et al. 1999). These structural changes are typically quantified at the plant and canopy level, where the impacts of N-limitation manifests over weeks to months, using optical or active LiDAR data (Berger et al. 2022). Whilst canopy structure and leaf biochemistry are relatively static in time, changes in LE can be detected more rapidly via thermal sensors at up to the minute scale (Vialet-Chabrand and Lawson 2020). When both optical and thermal sensors are combined, greater predictive modelling capacity is possible (Caine et al. 2024). This has the potential to optimise fertiliser application, enhance irrigation strategies, improve carbon flux modelling, and ultimately, boost overall yield forecasts.

The rapid development of new sensor technologies and their ongoing miniaturisation offers new opportunities to probe plant function in near-real time and at the individual plant scale (Guo et al. 2021; Xie and Yang 2020). Drone platforms have enabled within-field monitoring, at user-defined temporal intervals, transforming the observational scales at which it is possible to model crop performance (Guo et al. 2021; Messina and Modica 2020). However, different sensor technologies are often used in insolation and fail to harness the combined information content from different wavelength ranges that relate to time-dependent changes in plant fluxes and traits associated with changes in the surrounding environment. In this paper, we investigate the potential use of both optical and thermal-based remote sensing technologies to map crop responses in biochemical, structural and physiological traits, water fluxes and overall yields according to N fertiliser treatments with or without sucrose application in winter wheat. We address the following research questions: 1) Can N-imposed variations in crop water fluxes be captured using thermal data? 2) Can crop traits be mapped from optical drone data at the plant-scale? 3) Which crop traits and fluxes obtained from remotely-sensed data best serve as proxies for predicting crop yield?

## 2. Methods

### 2.1. Site description

Fieldwork was conducted on winter wheat (*Triticum aestivum* L.) variety Graham during a nutrient application field trial in East Yorkshire, UK (latitude: 54.0929, longitude: -0.6803) in June 2021. The average annual maximum temperature is 12.4°C and average annual rainfall is 771.16 mm, over a 1991-2020 climate period. For the field trials, there were 8 nutrient treatments in total, including a sucrose/no sucrose addition (Table 1). For the Regular N and Little and Often N (LO N) treatments, the same amount of total fertiliser was applied, but for the LO N treatment this was split into more frequent, smaller applications. The N fertiliser applied was Nitram (CF Fertilisers, UK). Sulphur was added to the Regular N treatment type (Regular NS) in the form of ammonium sulphate to promote N uptake. Each treatment plot was 2.1m x 24 m in size, and for each treatment there were three plot replicates.

**Table 1:** Details of the nutrient application for each treatment.

| Treatment | Nitrogen | Sulphur | Sucrose/ No Sucrose |
| --- | --- | --- | --- |
| 1. None | None | None | No |
| 2. Regular N | 290 kg/ha – 3 splits | None | No |
| 3. Little and often N<br>(LO N) | 290 kg/ha – 6 splits | None | No |
| 4. Regular N and S | 290 kg/ha – 3 splits | 50 kg/ha SO<sub>3</sub> | No |
| 5. None | None | None | Yes |
| 6. Regular N | 290 kg/ha – 3 splits | None | Yes |
| 7. Little and often N<br>(LO N) | 290 kg/ha – 6 splits | None | Yes |
| 8. Regular N and S | 290 kg/ha – 3 splits | 50 kg/ha SO<sub>3</sub> | Yes |

### 2.2. Ground measurements

#### 2.2.1. Leaf area index

Effective leaf area index (LAI) measurements were collected using the LAI-2200 plant canopy analyser (Li-Cor, Lincoln, NE, USA). A 45° disc was used to shield the instrument operator and reduce measurement uncertainty. For each plot, two above-canopy measurements were collected immediately before and after a sequence of 10 below-canopy measurements (Fang et al. 2019).

#### 2.2.2. Leaf gas exchange

Field measurements of leaf-level gas exchange were carried out using a LI-6800 portable infrared gas analyser (LI-COR, Lincoln, NE, USA). Flag leaves were used for all measurements including assessment of chlorophyll and nitrogen analysis. The LI-6800 was fitted with a 6400-02B Red/Blue Light Source, and CO_2_ response curves (A–C_i_ curves) were produced under light-saturating conditions, at photosynthetic photon flux density (PPFD) levels of 1800 μmol m^−2^ s^−1^, and stepwise CO_2_ concentrations of 420, 200, 100, 50, 25, 420, 420, 600, 800, 1000, 1200, 1500, 1800 μmol CO_2_ mol^−1^ air. Prior to logging measurements, leaves were acclimated in the chamber at 1800 μmol m^−2^ s^−1^, 60% relative humidity, a temperature of 25°C and a CO_2_ concentration of 420 μmol CO_2_ mol^−1^ air until steady-state conditions were reached. Throughout the measurement sequence, the leaf chamber was maintained as close to 25°C as possible (approximately ±1°C) and relative humidity was set to 60%. A complete *A*/*C*_i_ response curve took approximately 1 h to perform. The photosynthetic parameters *V*_cmax_ and *J*_max_, normalised to 25 °C, were calculated from the A-Ci curves fitted using the “Plantecophys” R package (Duursma 2015). Operating stomatal conductance measurements were collected between 11am and 1pm using a LI-600 porometer (Li-Cor, Lincoln, NE, USA) set to a flow rate of 150 μmol s^−1^. A total of 5 leaf measurements per plot were collected.

#### 2.2.3. Leaf biochemical analysis

Leaves were sampled, placed in open plastic bags and kept at a temperature of 0 °C during transport back to the laboratory for subsequent biochemical analysis to extract leaf chlorophyll and leaf nitrogen content. Foliar chlorophyll was extracted using spectra-analysed grade *N,N-*dimethylformamide, and absorbance was measured at 663.8, 646.8 and 480 nm using a UV-2600 spectrophotometer with an accompanying 6-cell CPS-100 cell positioner (Shimadzu, Kyoto, Japan). Leaf samples were dried at 80°C for 72 h and ground to a powder in a tissue lysis machine using metal 3mm ball bearings in 2 ml microcentrifuge tubes. Isotope analysis of % nitrogen content was obtained as in Field et al.(2016).

#### 2.2.4. Leaf hyperspectral reflectance and ground-based thermal imaging

Hyperspectral leaf reflectance from the adaxial surface of all sampled leaves was collected using a Spectral Evolution PSR+ 3500 spectroradiometer (Spectral Evolution, Inc. Massachusetts, USA), with a leaf clip attachment and internal calibrated light source. Spectra were automatically corrected for dark current and stray light, and referenced against an integrated Spectralon white reference standard (Labsphere, New Hampshire, USA). The PSR+ 3500 samples spectral radiance from 350-2500 nm across 2151 channels, for each measurement 25 spectra were internally averaged to increase the signal-to-noise ratio of the data.

A hand-held T650SC thermal imaging camera (FLIR Inc., Danderyd, Sweden) was used to collect thermal images of the sunlit top of canopy wheat plants for each sampled treatment plot. Measurements were collected during stable ‘blue sky conditions’ between 11am and 12pm on 09.06.21 as steady-state photosynthesis measurements were ongoing, at a time closest to that of the of the airborne thermal drone campaign (section 2.3.2). The T650SC camera operates in the 7.5–14 µm electromagnetic region with a 640 × 480-pixel line scan imager, providing a thermal sensitivity of < 20 mK. The thermal images were taken at a constant height of c. 1.8 m and a distance of 1 m from the plant. Each thermal image approximately covered an area of approx. 0.70 × 0.70 m.

#### 2.2.5. Hyperspectral and partial least squares regression data analysis

Hyperspectral datasets were imported into R via the Spectrolab package and partial least squares regression analysis was conducted using the Spectratrait package (Burnett et al. 2021; Meireles et al. 2017). To build a model to predict leaf chlorophyll content 116 samples were analysed: 92 for the calibration dataset (79%) and 24 for validation dataset (21%).

#### 2.2.6. Leaf stomatal analysis

Epidermal imaging assessment was undertaken on ImageJ using nail varnish impressions of dental resin moulds taken from flag leaves of plants growing in the respective treatments. Two 0.44 mm^2^ fields of view (FOV) approximately 8 veins from the leaf margin were assessed and then averaged to compute stomatal size and stomatal density for each replicate. For stomatal size calculation, 5 individual guard cells lengths were measured, then average per FOV.

### 2.3. Drone data acquisition

#### 2.3.1 Optical drone measurements

Airborne multispectral data was collected by a DJI Matrice M200 quadcopter (Nanshan, Shenzhen, China) equipped with MicaSense RedEdge-Mx multispectral imaging sensor (MicaSense, Seattle, WA, USA) between 11.00 and 12.00 GMT on June 6th, 2021. The RedEdge-Mx sensor acquires images in 5 different spectral bands: blue (475 nm centre, 32 nm bandwidth), green (560 nm centre, 27 nm bandwidth), red (668 nm centre, 14 nm bandwidth), red edge (717 nm centre, 12 nm bandwidth) and near-infrared (840 nm centre, 57 nm bandwidth). The flight was flown an altitude of 10 m above the ground, giving a spatial resolution of 0.52 cm. The airborne surveys consisted of 994 images. The images were radiometrically calibrated using a calibrated reference panel (MicaSense, Seattle, WA, USA) captured immediately before and after each flight, and was accompanied by an additional DLS 2 Downwelling Light Sensor which determines the ambient light and sun angle for each band.

#### 2.3.2. Thermal drone measurements

Land surface temperature data was sampled using a FLIR Lepton 3.5 microbolometer radiometric thermal sensor on board a Parrot Analfi drone (Paris, France). The flight was performed between 10.00 and 11.00 GMT on June 9^th^, 2021 at an altitude of 10 m above the ground, giving a spatial resolution of 0.34 cm. The Lepton 3.5 sensor acquires images in the 8–14 µm range with a 160 x 120 pixel sensor array, and a focal length of 26 mm, giving a horizontal field of view of 57°. This sensor has a thermal sensitivity of <50mK. Flights were carried out with an 80% forward overlap and 70% side overlap. The airborne surveys consisted of 473 images. Simultaneous Aerial RGB Images were also collected via a Sony CMOS HDR sensor 1/2.4’’ 21MP digital camera (Tokyo, Japan) on board the Parrot Anafi drone.

#### 2.3.3. Drone data processing

The optical and thermal imaging data was processed in Metashape (Agisoft, St. Petersburg, Russia) to create geometrically corrected orthomosaics. For thermal orthomosaic construction, individual thermal photograph temperatures were first standardised in FLIR Researcher IR software. A total of 10 spot measurements per plot were collected to produce the average plant temperature per plot. The default settings in PIX4Dmapper (Pix4d, Lausanne, Switzerland) were used to align and produce the RGB orthomosaic presented in Fig. 1.

**Fig. 1:**
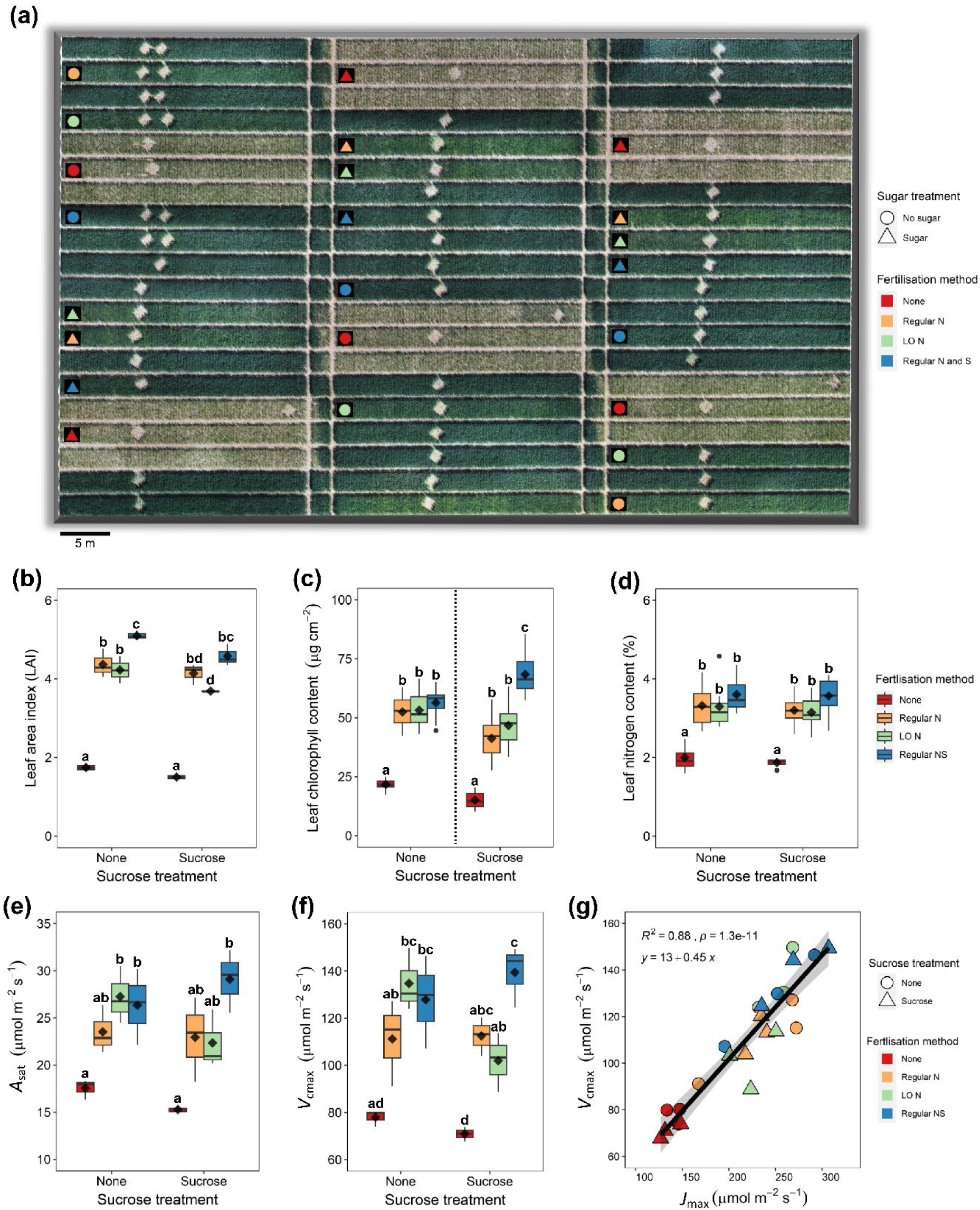
Overview of multi-treatment wheat nutrient experiment with ground-based trait measurements. (**a**), Drone derived RGB orthomosaic highlighting the 24 plots studied with individual treatments marked with different symbols and colours (see key). Ground measurements corresponding to: (**b**), leaf area index (LAI; *n* = 3); (**c**), leaf chlorophyll content (*n* = 14 or 15); (**d**), leaf nitrogen content (%; *n* = 8); (**e**), photosynthesis (*A*_sat_; *n* = 3) and (**f**), Maximum carboxylation rate (*V*_cmax_; *n* = 3). (**g**), Regression between *V*_cmax_ and maximum rate of electron transport (*J*_max_; *n* = 3). Traits in Figs **b**, **d**, **e** and **f** were assessed using Two-way ANOVAs, with Tukey post-hoc tests. In **c**, a dashed line indicates different sucrose treatment samples were assessed individually using two separate Welch’s ANOVAs, followed by Games-Howell post-hoc tests. Boxplot whiskers indicate the minimum and maximum values, black dots represent outliers, straight horizontal lines represent medians and diamonds represent samples means. Different letters denote *P* < 0.05 or higher significance.

### 2.4. Yield data collection

The grain yield of all plots was recorded using a Sampo small plot combine harvester (Sampo Rosenlew Ltd., Pori, Finland) with a vertical cutting bar, cutting the full plot width of 2.1m. Areas of the plot from which quadrat samples were collected were excluded from the harvested area. Harvested plot lengths were measured immediately after harvest and grain yield adjusted for both plot size and moisture content. The moisture content of the grain was determined using a Dickey John GAC 2000 grain analysis computer. The grain yield was adjusted to a standard moisture content of 15%.

## 3. Results

### 3.1. Impacts of N-fertiliser and sucrose application on wheat traits and fluxes

To assess crop responses to different N-fertiliser treatment regimes, we assessed a comprehensive range of plant structural, biochemical, physiological and spectral traits (Figs. 1-2). Clear differences between fertilised (N) and unfertilised (None) plants can be seen in a drone-based RGB orthomosaic, with Regular NS plants (that received additional S) typically having darker green canopies (Fig. 1a).

**Fig. 2:**
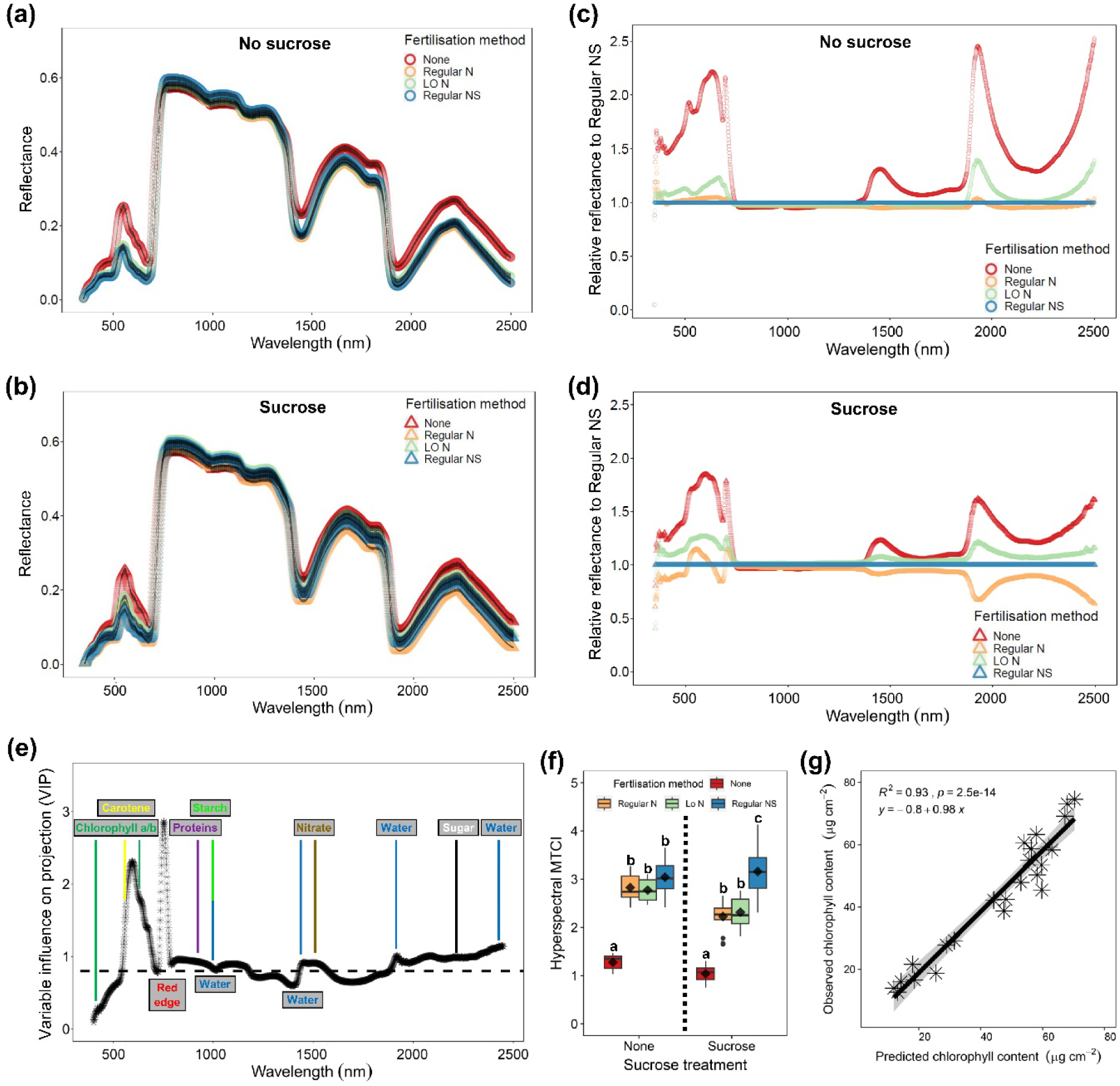
Hyperspectral leaf analysis of wheat grown under differing fertiliser and sugar applications. Hyperspectral reflectance profiles of (**a**), four fertiliser treatments grown in the absence of sucrose and (**b**), the same treatments grown with sucrose. Relative reflectance comparatively to Regular NS treatment for (**c**), no sucrose plants, and (**d**), sucrose treated plants. (**e**), Variable Influence of Projection (VIP) graph from partial least squares regression (PLSR) analysis of chlorophyll from the 8 treatment groups presented in **a** and **b**, along with known absorption features associated with key cellular and molecular components (Au - Kuska et al. 2018). (**f**), Hyperspectral MERIS Terrestrial Chlorophyll Index (MTCI) values from different plots. (**g**), PLSR modelled predictions of leaf chlorophyll content. The dark markings in **a** and **b** associated with each sample are error bars equating to +/- 1 standard error of the mean. Boxplot whiskers indicate the minimum and maximum values, black dots represent outliers, straight horizontal lines represent medians and diamonds represent samples means. Different letters denote *P* < 0.05 or higher significance. Sample *n* = 14 or 15.

For all measurements collected, the absence of N resulted in the lowest trait values (Figs. 1b-g). N applied conventionally, in three larger applications (Regular N; methods Table 1), or applied ‘little and often’ (LO N), within each sucrose treatment (sucrose or no sucrose), did not significantly impact any traits measured in Fig. 1. Sucrose overall had a negative impact on leaf area index (LAI) but not on any of the other measured traits (Fig. 1b, Supplementary Table 1). LAI was also impacted by S application, with Regular NS plants having the highest or equal highest values within both sucrose treatments (Fig. 1b; *P* < 0.05, Two-way ANOVA, post-hoc TUKEY HSD test, Supplementary Table 1). Leaf chlorophyll was also highest or equal highest in Regular NS plants compared to Regular N (Fig. 1c; *P* < 0.001, Welches ANOVA, post-hoc Games-Howell test, Supplementary Table 1), but no significant differences were detected between any of the N-treated plants (including Regular NS) for leaf N content or light-saturated photosynthesis (*A*_sat_; Fig. 1d and e). Like chlorophyll content, *V*_cmax_ values also revealed Regular NS to be beneficial to performance, particularly in the presence of sucrose (Fig. 1f; *P* < 0.05, Two-way ANOVA, post-hoc TUKEY HSD test, Supplementary Table 1). Overall N allocation strategy between electron transfer (*J*_max_) and *V*_cmax_ remained consistent with N treatments supplied either with or without sucrose (Fig. 1g; *R^2^* = 0.88, *P* < 0.001). No significant differences in photosynthetic nitrogen-use efficiency (photosynthesis/nitrogen content; pNUE) or intrinsic water-use efficiency (*A*_sat_/stomatal conductance (*g*_sat_); iWUE) were detected between any of the treatments (Supplementary Fig. 1, Supplementary Table 1).

Hyperspectral reflectance data collected from the same leaves as in Fig. 1, are presented in Fig. 2a-b. Figs. 2c and 2d show that LO N plants typically absorb less radiation than Regular N plants across much of the electromagnetic spectrum (particularly in the visible spectrum and at 1450 and 1920 nm wavelengths), indicative of lower leaf pigment and leaf water content. Sucrose altered the overall reflectance properties of fertilised plants relative to unfertilised equivalents by reducing overall differences (particularly between 1920-2500 nm) (Fig. 2b and d). Partial least squares regression analysis (PLSR) against measured leaf chlorophyll content confirmed the red-edge as the most important wavelength range for quantifying the variation in observed chlorophyll content within Variable Influence on Projection (VIP) plots (Fig. 2e). Four regions in the near infrared (NIR) and shortwave infrared (SWIR) often associated with different water bands (970, 1450, 1900 and 2500 nm) were also important contributors for predicting chlorophyll content, as was a nitrate band at 1500 nm and a sucrose band around 2250 nm (Au - Kuska et al. 2018). Calculation of the chlorophyll-sensitive Meris Terrestrial Chlorophyll Index (MTCI) revealed Regular NS had the highest or equal highest values (Fig. 2f; *P* < 0.0001, Welches ANOVA, post-hoc Games-Howell test). Finally, PLSR modelling provided a very accurate prediction of chlorophyll content across different fertiliser and sucrose treatment combinations (Fig. 2f; *R^2^* = 0.93, *P* < 0.001).

The different N-based fertiliser and sucrose treatments applied resulted in changes to both the biochemistry and physiology of wheat leaves (Figs. 1 and 2). To understand how plants responded at the cellular level to shifts in demands for CO_2_ substrate (and associated water loss changes), we measured leaf stomatal traits and found that N-fertilised plants had significantly larger stomata size (SS) and often displayed a trend towards lower stomatal density (SD; Fig. 3a-f, Supplementary Table 2). SD appeared linked to stomatal files per mm^-2^ (Fig. 3g), as N fertilisation (either regular N, LO N, or regular NS) produced stomatal files further apart. Sucrose reduced SS, and increased SD and stomatal final number (Fig 3, Supplementary Table 2). Porometry assessment of stomatal conductance (*g_sw_*) revealed all 3 N-fertilised treatments had significantly higher values than no fertiliser wheat, but overall sucrose treatment did not alter *g_sw_* (Fig. 3h, Supplementary Table 2).

**Fig. 3.**
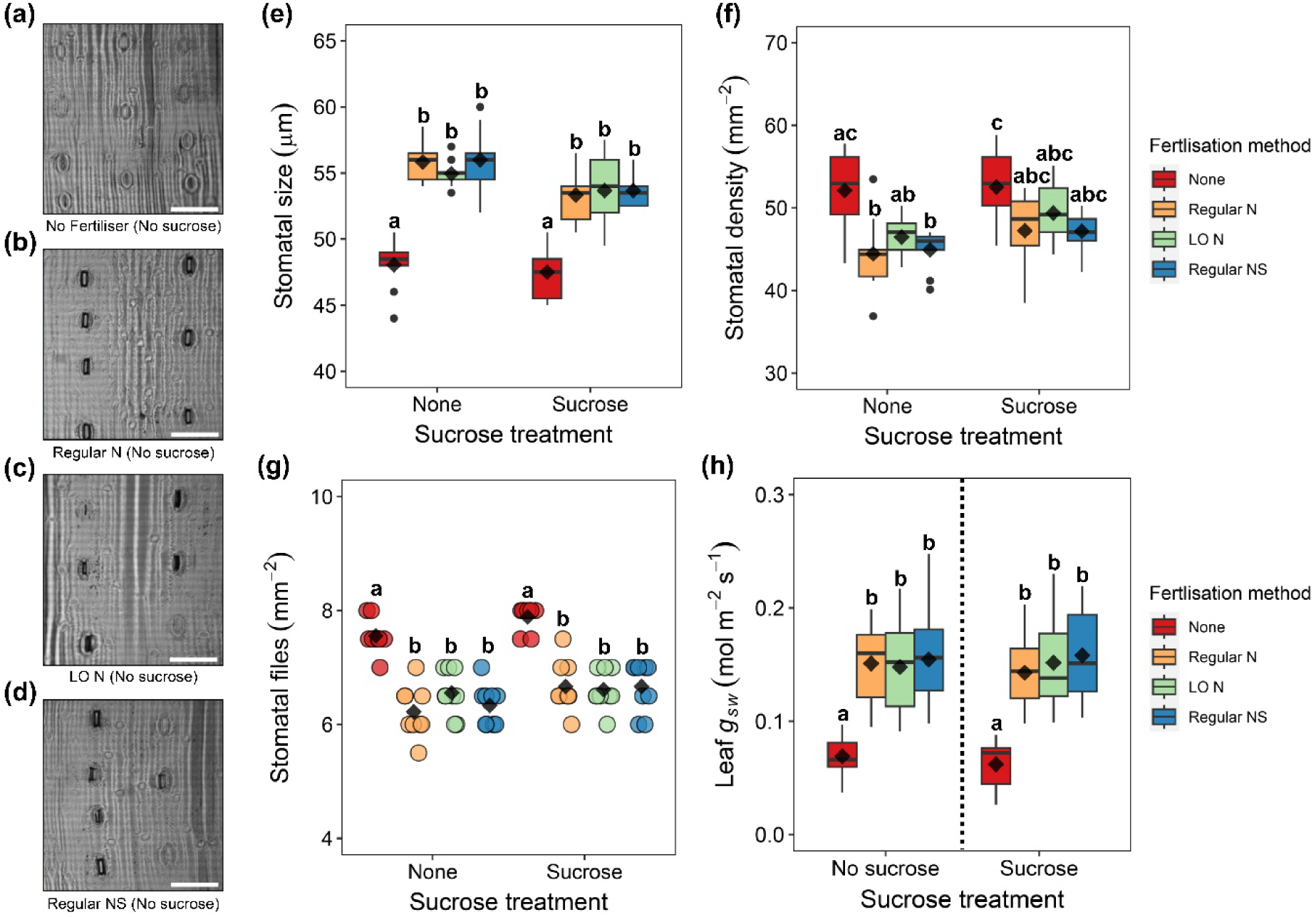
Wheat leaf stomatal properties and corresponding gas exchange. a-d, Representative stomatal images of (**a**), No fertiliser (None), (**b**), Regular N, (**c**), LO N and (**d**), Regular NS treatments with no additional sucrose. (**e**), Stomatal size, (**f**), stomatal density, (**g**), stomatal file number and (**h**), operating stomatal conductance (*g_sw_*) of plants treated with different fertiliser and or sucrose. For **e**-**g**, *n* = 9, for **h**, *n* = 15. Two-way ANOVAs were conducted for each trait, with post-hoc Tukey tests to compute significance. Boxplot whiskers indicate the minimum and maximum values, black dots represent outliers, straight horizontal lines represent medians and diamonds represent samples means. In **h**, a dashed line indicates different sucrose treatment samples were assessed individually using two separate Welch’s ANOVAs, followed by Games-Howell post-hoc tests. Different letters imply *P* < 0.05 or higher significance. Scale bars in A-D = 100 µm.

### 3.2. Capturing nutrient-induced variations in water fluxes from thermal imagery

To investigate if thermal RS can accurately quantify differences in *g_sw_* between different fertiliser and sucrose treatments, plant leaf temperatures were sampled using drone-based thermal imagery, alongside ground-based images for comparison (Fig. 4).

**Fig. 4.**
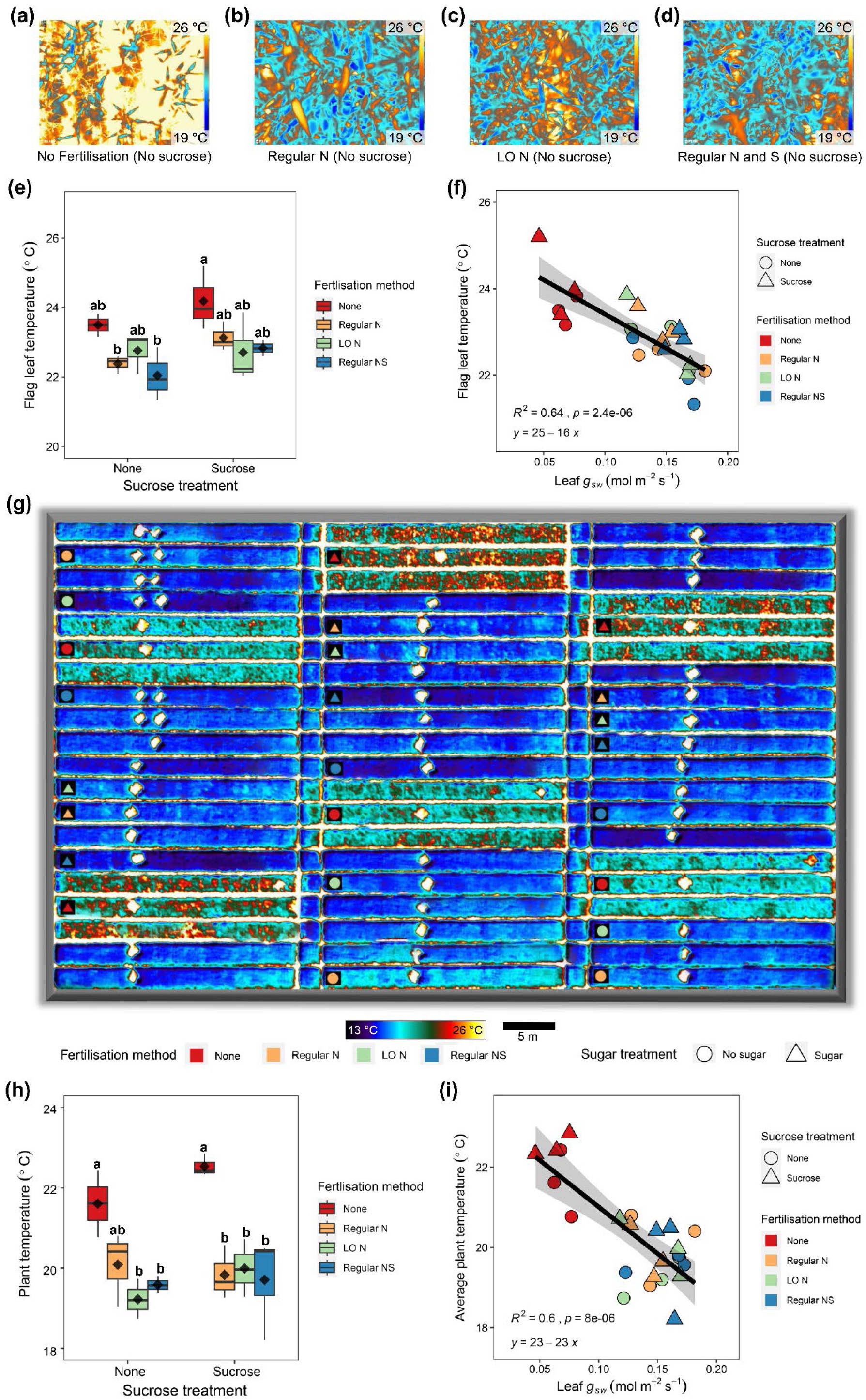
Ground and aerial based thermal imaging of wheat plants grown using different Nitrogen (N) fertiliser, Sulphur (S) and or sucrose treatments. Representative ground thermal images (**a**), No N fertiliser (None), (**b**), Regular N, (**c**), Little an often N (LO N) and (**d**), Regular NS. (**e**), Individual flag leaf temperatures (*n* = 9). (**f**), Plot averaged regression analysis of flag leaf temperature vs stomatal conductance (g_sw_) (*n* = 4 or 5). (**g**), Thermal orthomosaic from drone thermal imagery of all 24 plots surveyed in the study. (**h**), Temperatures of plants of different treatments (10 spot measurements per plot). **i**, Regression analyses investigating the relationships between leaf temperature and ground-collected g_sw_ measurements at the plot-level. In **e** and **h**, Two-way ANOVAs with Tukey post-hoc tests were conducted. Boxplot whiskers indicate the minimum and maximum values, straight horizontal lines represent medians and diamonds represent samples means. Different letters denote *P* < 0.05 or higher significance. Scale in **g** = 5 m.

Results of ground-based thermal imaging revealed that all forms of N-fertiliser application increased plant cooling (Fig. 4a-e; Two-way ANOVA, *P* < 0.05) but sucrose did not have a significant impact. Leaf-level regression analyses revealed flag leaf temperature (*R^2^* = 0.64, *P* < 0.001) to be a good indicator of *g_sw_* (Fig. 4f). The thermal orthomosaic revealed that unfertilised plants were significantly warmer than fertilised equivalents (Fig. 4g-h). There were no statistical differences between the temperatures of plants that received N fertiliser (including Regular NS treatment), and overall sucrose did not have a significant impact (Fig. 4h). Regression analysis revealed that the average plant temperature per plot (Fig. 4i; *R^2^*= 0.6, *P* < 0.001) served as a good proxy for ground-based *g_sw_*measurements.

### 3.3. Mapping nutrient-induced variations in physiological traits from drone-based optical imagery

Previous work has shown that leaf chlorophyll is a strong proxy for *V*_cmax_ (Croft et al. 2017). To produce spatially-continuous maps of *V*_cmax_ over the different treatment plots, we used the MTCI spectral vegetation index (Fig. 5).

**Fig. 5:**
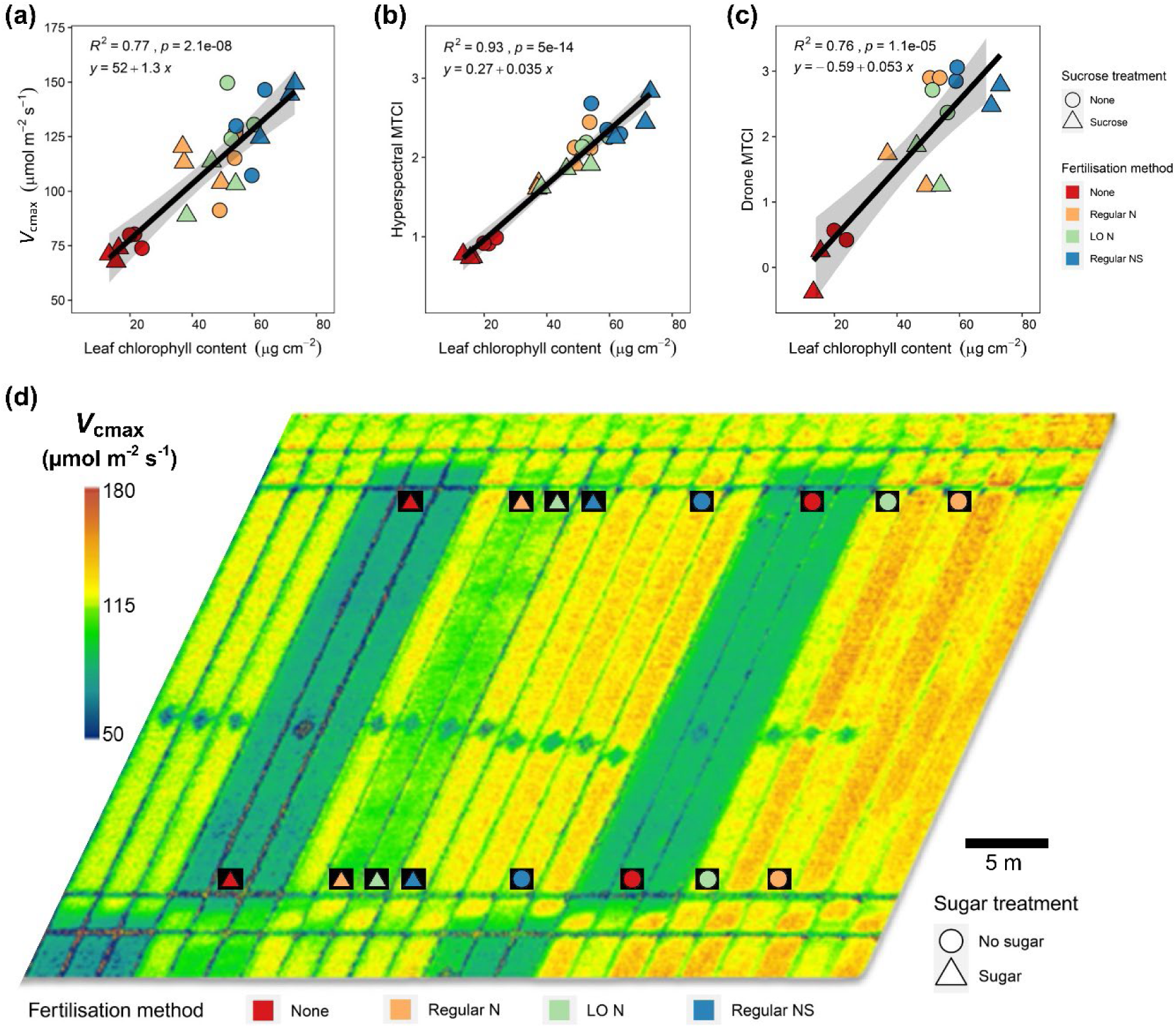
Mapping of maximum rate of rubisco carboxylation (*V*_cmax_) to wheat canopies using drone-derived MERIS Terrestrial Chlorophyll Index (MTCI). (**a**-**c**), Plot level regression analyses of measured leaf chlorophyll content (*n* = 4 to 5 per plot) with (**a**), *V*_cmax_ (*n* = 1 per plot), (**b**), Hyperspectral MTCI (*n* = 4-5 per plot) and (**c**), Drone MTCI (*n* = 3 per plot). (**d**), Mapped V_cmax_ based on drone MTCI. Scale in D = 5 m.

Leaf-level measurements of *V*_cmax_ presented a strong relationship with leaf chlorophyll content (*R^2^* = 0.77, *P* < 0.001) regardless of treatment (Fig 5a). Level-level MTCI and drone-based MTCI values both correlated well with measured leaf chlorophyll content (*R^2^* = 0.93, *P* < 0.001 and *R^2^* = 0.76, *P* < 0.001), respectively (Figs. 5b-c), providing an optical basis for scaling *V*_cmax_ across space. Mapped *V*_cmax_ reveals similar productivity responses between different fertiliser and sucrose treatments to those found with ground-based measurements (Fig. 5d, see also Fig. 1f). Regular NS application led to the most productive (or equal most productive) plots, whereas sucrose application often reduced productivity, and no fertiliser application resulted in the lowest *V*_cmax_ values.

### 3.4. Spectral, trait and flux relationships with final grain yield

The efficacy of using peak season drone data for predicting N-fertiliser induced variations in crop yield, grain yields from all plots were quantified at the end of the season is investigated in Fig. 6a-b. Regular NS plants had the highest or equal highest yields, producing c. 11.5-12.3 tonnes per hectare (Fig. 6c). Compared to Regular N, Regular NS had increased yields of 13% over the no sucrose treatment and 24% over the sucrose treatment. Overall sucrose had a strong negative impact on yields (Two-way ANOVA, *P <* 0.001), but no difference was detected in rate of N application (Regular N vs LO N) within each sucrose treatment.

**Fig. 6.**
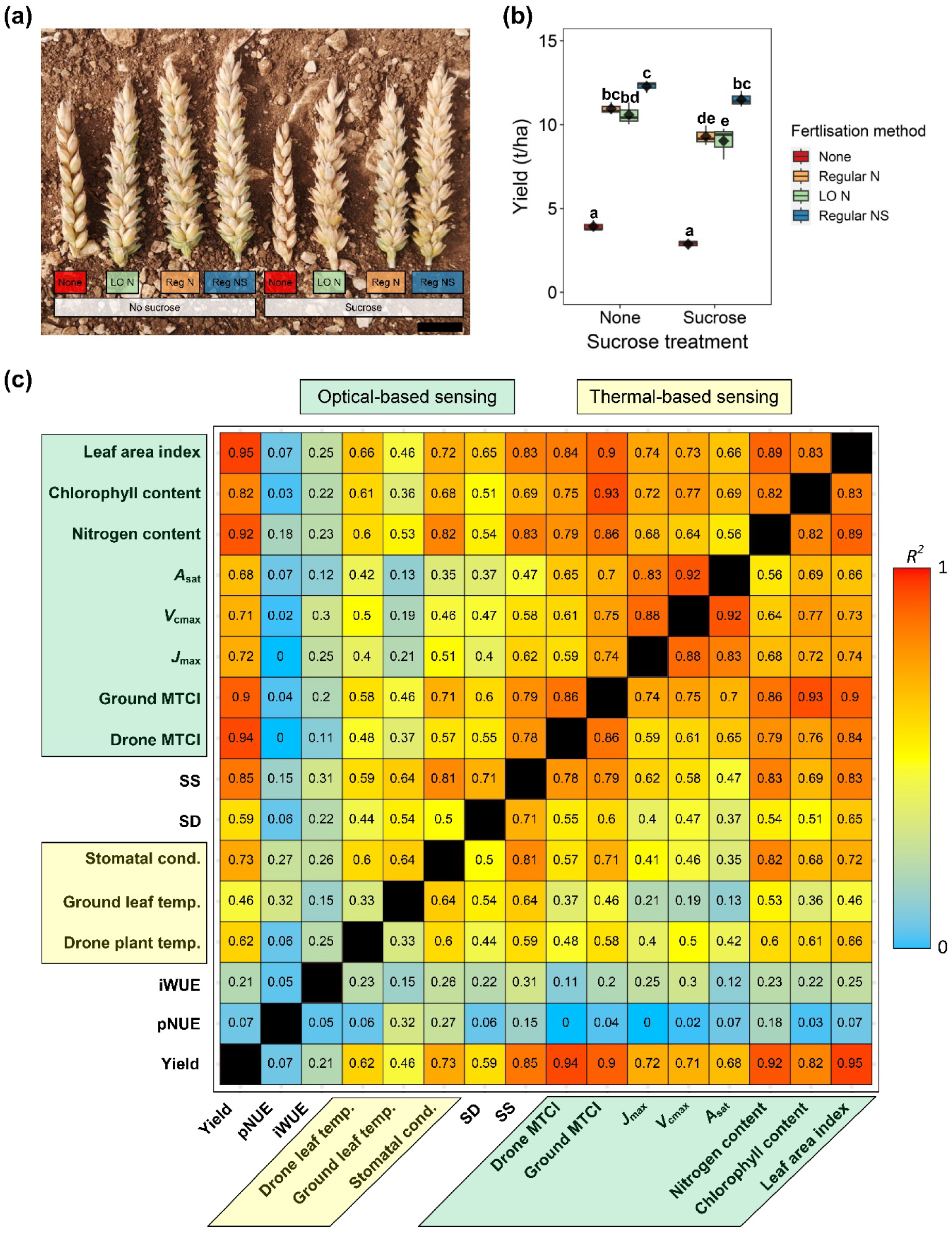
Yield quantification and relationship to measured plant traits and fluxes at the canopy level. (**a**), Examples of wheat ears. (**b**), Quantification of yield (*n* = 3 plots per treatment). (**c**), Correlation matrix of measurements at the plot level highlighting R^2^ relationships of yield with other plant traits and fluxes (*n* = 2 or 3 plots per treatment). Traits associated with optical-based sensing have green backgrounds and fluxes that are associated with thermal-based sensing have light yellow backgrounds. Darker orange to red squares with the matrix indicate stronger correlations.

Plot-level correlation matrices show relationships between yield, physiological parameters and optical and thermal data (Fig. 6c, Supplementary Fig. 2). Very strong positive relationships were found between yield and LAI (*R^2^* = 0.95), drone MTCI (*R^2^* = 0.94) and hyperspectral ground MTCI (*R^2^* = 0.90). Yield was also very strongly associated with leaf nitrogen (*R^2^* = 0.92) and chlorophyll content (*R^2^* = 0.82), and to a lesser extent with *J*_max_ (*R^2^* = 0.72), *V_cmax_*(*R^2^* = 0.71) and *A*_sat_ (*R^2^* = 0.68). Plot measurements of pNUE and iWUE only showed limited or no relationships with other plant traits, but both SS (*R^2^* = 0.85) and SD (*R^2^* = 0.58) showed noticeable relationships with yield, with larger SS lower SD plants typically producing higher yields. Assessment of yield with thermally acquired flux data including drone plant temperature (*R^2^* = 0.62) and ground leaf temperature (*R^2^* = 0.46) showed good to moderate relationships, with cooler plants typically having higher yield. Confidence in thermal-based flux estimates is further underpinned by a robust relationship between leaf *g_sw_* and overall grain yield (*R^2^*= 0.73).

## 4. Discussion

### 4.1. Modelling dynamic responses in plant fluxes

Under limited N, crops adjust their physiological, biochemical and structural traits (Fig. 1; Section 1, Section 3.1). The nature, extent and timings of these adjustments has previously been shown to vary according to the severity and duration of the stress (Berger et al. 2020). As N first becomes limiting, changes primarily occur in the fluxes of water, carbon and energy exchanged between plants and the atmosphere. These early changes in fluxes therefore offer an ‘early-warning’ for crop management before crop changes become visible and are yield-limiting. In the short term, this occurs via stomatal pore aperture changes caused via adjustments to guard cell turgor and over longer durations, plants can alter the SS and SD of stomata in response to changes in the environment (Bertolino et al. 2019). Here, we found that increased N availability increased SS and reduced SD, and increased *g_sw_* of water vapour (Fig. 3 and 4; Sections 3.1 and 3.2). Thermal data was able to detect these stomatal level responses within a space-for-time framework, presenting a strong relationship with stomatal conductance (Figure 4, 6). This strong inverse relationship found between leaf temperature and *g_sw_* is due to evaporative cooling as plants transpire water through stomata (Jones 2004). Previous studies have shown the relationship between water fluxes at the stomatal scale and thermal imagery in laboratory or controlled environment experiments (Caine et al. 2023; Leinonen et al. 2006; Vialet-Chabrand and Lawson 2019). However, few have demonstrated the potential in situ drone-based thermal imagery, for capturing fluxes at the stomatal rather than canopy scale. A notable exception includes Brewer et al., (2022), who used thermal UAV data to model stomatal conductance in maize with good accuracy (between *R^2^* = 0.58 and *R^2^* = 0.74; for different phenological stages).

### 4.2. Modelling trait-based responses to N limitation

Under more severe N limitation, leaf chlorophyll content Rubisco content, bioenergetics and light-harvesting proteins decreased, reducing electron transport rate and carboxylation, and ultimately photosynthesis (Figure 1) supporting the findings in (Caine et al. 2024; Mu and Chen 2021). These changes led to lower sugar production due to reduced photosynthesis, which decreases growth rates leading to smaller plant stature (Berger et al. 2020; Caine et al. 2024). Through a space-for-time perspective, the findings here capture the impact of N limitation and demonstrate their detection at different stages of N stress using optical-based techniques (Fig. 1-6). At the leaf level, clear differences were seen in the hyperspectral reflectance between the treatments, largely in spectral regions associated with relating to pigment and water absorption bands (Fig. 2). PLSR analysis enabled the accurate modelling of leaf chlorophyll content from hyperspectral data (*R^2^* = 0.93; *P* < 0.001), supporting the efficacy of this data-driven technique in modelling differences in plant traits shown in other studies (Burnett et al. 2021). When scaling to spatially-continuous image-based reflectance data from drones, the availability of hyperspectral data is often restricted due to prohibitive costs, leading to a trade-off between the spatial and spectral domains. Drone-mounted (and satellite) sensors are often multispectral. In this case, we found the MTCI spectral vegetation index presented strong relationships with leaf chlorophyll and LAI (*R^2^* = 0.76; *P* < 0.001 and *R^2^* = 0.84; *P* < 0.001, respectively). The close relationship between chlorophyll content and Vcmax (Croft et al., 2019) allowed the mapping of photosynthetic capacity at the individual plant scale, revealing variations in the field both between and within the treatments, and offering the potential for precise and targeted management techniques.

### 4.3. Modelling longer-term crop acclimation to N-fertiliser schemes and final yields

Whilst research has shown that sucrose may promote beneficial plant-microbe interactions (Yang et al. 2022), the results here found that sucrose addition often hindered plant performance. Combined N and S application produced the largest, or equal largest, final grain yields, which was underpinned by enhanced biochemical and physiological performance during the growing season (Figs. 1-6). Sulphur (S) is also a key element for optimal wheat growth, and is essential for producing a number of amino acids, proteins and for chlorophyll biosynthesis (Jamal et al. 2010), and additionally promotes N uptake (Salvagiotti et al. 2009). Our data aligns with Fatma et al. (2023) who showed combined N and S produced the highest trait values for several biochemical and physiological traits, including chlorophyll content, photosynthesis, rubisco activity and the electron transport rate. In other crop species, research found that under N limitation, the fraction of N supplied to the electron transport chain increased by 17%, and the fraction of N invested in carboxylation declined by 54% (Zhong et al. 2019). This suggests that under nitrogen stress, a greater N investment into capturing light energy than fixing carbon via Rubisco (Mu and Chen 2021; Zhong et al. 2019). However, we found a strong linear relationship between *V*_cmax_ and *J*_max_ (Fig. 1) regardless of N-fertiliser (and or sucrose) treatment, implying no adjustment in the allocation of N to bioenergetics in this case.

Structural and biochemical traits (and their optical proxies) reflect a longer-term integrated signal of growing season conditions, and therefore an extremely strong statistical relationship with yield (*R^2^*= 0.94; *P* < 0.001, for drone-based MTCI). The strong relationship between MTCI and leaf chlorophyll and LAI (Section 4.2, Figure 6) indicates that MTCI represents the likely fraction of absorbed PAR that would be used for photosynthesis (Zhang et al. 2009) and explains the strong relationship with yield. In partial contrast, variations in gas exchange (and its thermal proxy) reflect the physiological activity of the crop at that time, which may also be in response to diurnal variations and light dynamics and to meteorological conditions. Thermal data are thus dynamic and represent a snapshot of plant fluxes in time. As a consequence, the thermal data does not show as strong a relationship with final yield as optical remote sensing data, because of this real-time variability in the thermal signal. Nonetheless, thermal RS data could provide a dynamic stress constraint within a modelling framework. The integration of thermal data with meteorological and optical data to model the cumulative biochemical and structural response of crops to environmental drivers may provide important advances in the accuracy of yield forecasts in real time and the dynamic adjustment of fertiliser application.

## 5. Conclusion

Our results show that variation in N, S and/or sucrose all contributed to phenotypic differences in winter wheat, which were accurately retrievable from both optical and thermal drone imagery. The ability to spatially-map crop physiological adjustments to nutrient supply in traits and fluxes will revolutionise the precise management of crop nutrient needs to sustainably enhance overall yields. The application of different nutrient treatments found that S boosted wheat physiological performance and yields, whereas N application rate and timing had little overall effect and sucrose typically had a negative impact. Drone-based data showed strong relationships with corresponding ground-based traits and fluxes, with optical drone data showing the strongest potential for predicting grain yields (Sections 3.2-3.4). This research shows the power of combined optical and thermal drone phenotyping in an agricultural field setting and paves the way for developing sophisticated drone-informed models that enable targeted fertilisation strategies that reduce N emissions and maximise yields.

## Acknowledgements

This work was supported by the UK Research and Innovation (UKRI) Future Leaders Fellowship scheme [MR/T01993X/1] and The Institute of Sustainable Food at the University of Sheffield. The trial was funded by CF Fertilisers UK Ltd

## Author contributions

PMB, KES and HLC designed the study. BC, PMB, KES and HLC collected the data. BC and HLC undertook data analysis. BC and HLC wrote the manuscript with input from PMB and KES.

## Data availability

Please contact the corresponding author of this study should any data from the study be required.

