## Supplementary material for "Mapping plant-scale variation in crop physiological traits and water fluxes"

**Supplementary information**

Supplementary Table 1. Statistical analysis overview of results presented in Fig. 1. All traits except chlorophyll content were assessed using Two-way ANOVAs with Tukey post-hoc tests. Chlorophyll content of no sucrose and sucrose treated samples were assessed using individual Welch ANOVA tests. Regular N vs little and often N (LO N) and Regular N vs Regular N and S (Regular NS) tests are aggregated for sucrose and no sucrose treatments. Significance is as follows; $P < 0.05$ = *, $P < 0.01$ = **, $P < 0.001$ or greater significance = ***. NS means not significant.

| Plant trait measurement | Fertiliser significance | Sucrose treatment significance | Treatment interaction | Aggregated Regular N vs LO N | Aggregated Regular N vs Regular N and S |
|---|---|---|---|---|---|
| Leaf area index (LAI) | *** | *** | NS | NS | ** |
| % Nitrogen content | *** | NS | NS | NS | NS |
| Photosynthesis | *** | NS | NS | NS | NS |
| Max. rate of rubisco carboxylation ($V_{cmax}$) | *** | NS | * | NS | * |
| Chlorophyll content Sugar treatment (None) | *** | NA | NA | NA | NA |
| Chlorophyll content Sugar treatment (Sugar) | *** | NA | NA | NA | NA |
| Nitrogen-use efficiency (pNUE) | NS | NS | NS | NA | NA |

Supplementary Table 2. Statistical analysis overview of results presented in Fig. 3. All traits except stomatal conductance were assessed using Two-way ANOVAs with Tukey post-hoc tests. Stomatal conductance measurements of no sucrose and sucrose treated samples were assessed using individual Welch ANOVA tests. Significance is as follows; $P < 0.05$ = *, $P < 0.01$ = **, $P < 0.001$ or greater significance = ***. NS means not significant.

| Plant trait measurement | Fertiliser significance | Sugar treatment significance | Treatment interaction | Aggregated Regular N vs LO N | Aggregated Regular N vs Regular N and S |
|---|---|---|---|---|---|
| Stomatal size | *** | ** | NS | NS | NS |
| Stomatal density | *** | * | NS | NS | NS |
| Stomatal file. no | *** | ** | NS | NS | NS |
| Stomatal conductance (none) | *** | NA | NA | NA | NA |
| Stomatal conductance (sugar) | *** | NA | NA | NA | NA |

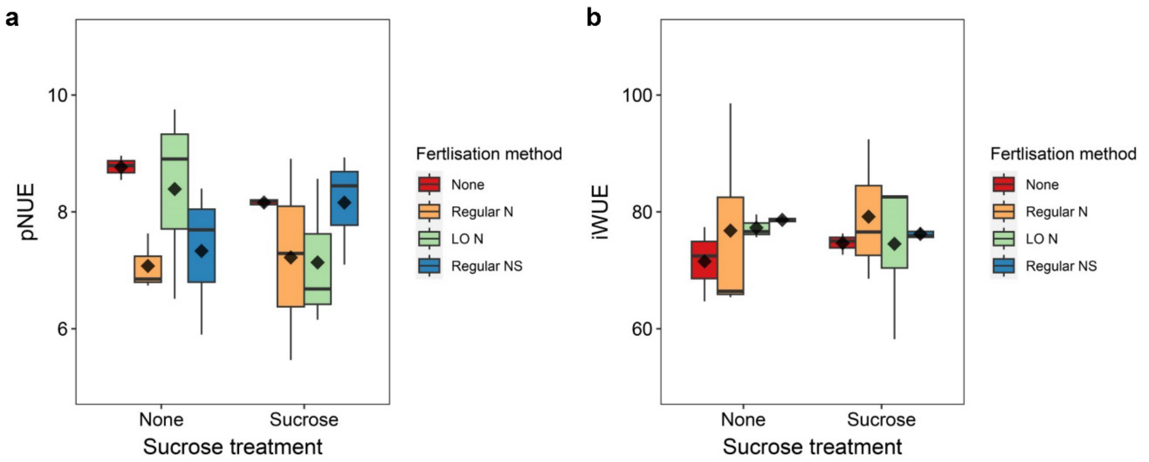


Supplementary Fig. 1. (a) Photosynthetic nitrogen-use efficiency and (b) intrinsic water-use efficiency of plants exposed to different fertilisation and or sugar treatment methods. Two-way ANOVAs were undertaken for both measured traits but no significant differences were detected. *n* = 3. Boxplot whiskers indicate the minimum and maximum values, black dots represent outliers, straight horizontal lines represent medians and diamonds represent samples means.

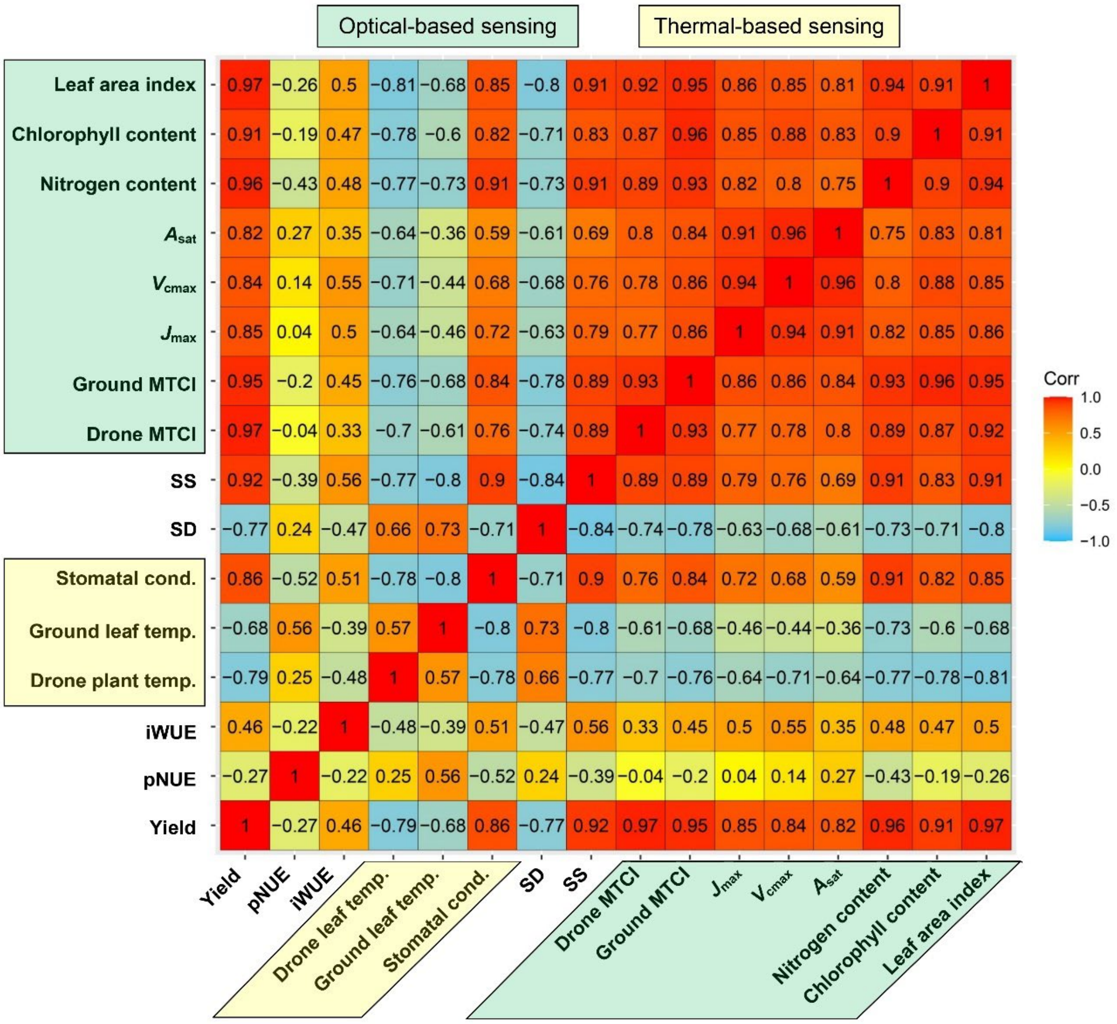


Supplementary Fig. 2. Regression co-efficient analyses of traits and fluxes present in Fig. 6. Orange to red colours represent strong positive relationships and lighter blue to sky blue colours represent strong inverse correlations.
